# An unusual two-strain cholera outbreak in Lebanon, 2022-2023: a genomic epidemiology study

**DOI:** 10.1101/2023.11.29.569232

**Authors:** Antoine Abou Fayad, Rayane Rafei, Elisabeth Njamkepo, Jana Ezzeddine, Hadi Hussein, Solara Sinno, Jose-Rita Gerges, Sara Barada, Ahmad Sleiman, Moubadda Assi, Maryo Baakliny, Lama Hamedeh, Rami Mahfouz, Fouad Dabboussi, Rita Feghali, Zeina Mohsen, Alisar Rady, Nada Ghosn, Firas Abiad, Abdinasir Abubakar, Amal Barakat, Nadia Wauquier, Marie-Laure Quilici, François-Xavier Weill, Monzer Hamze, Ghassan M. Matar

## Abstract

**Background:** Cholera is a bacterial infection caused by the ingestion of contaminated water or food. It principally affects the gastrointestinal system and spreads easily, causing outbreaks. The first case of cholera in this outbreak was detected in Lebanon in October 2022. The outbreak lasted three months, with 8,007 suspected cases (671 laboratory-confirmed) and 23 deaths. We characterised the *Vibrio cholerae* strain responsible for this cholera outbreak.

**Methods:** In total, 34 *Vibrio cholerae* isolates collected by random sampling of stools, water and plant samples throughout the outbreak and over the affected regions were studied by phenotypic methods and microbial genomics.

**Findings:** All isolates were *V. cholerae* O1, serotype Ogawa strains from wave 3 of the seventh pandemic El Tor (7PET) lineage. Phylogenomic analysis unexpectedly revealed the presence of two different 7PET strains, a highly unusual finding outside the Bay of Bengal, where several sublineages circulate together. The dominant strain had a narrow antibiotic resistance profile and was phylogenetically related to South Asian *V. cholerae* isolates. The second strain, which was found exclusively in South Lebanon and Beqaa, was resistant to multiple antibiotics, including macrolides, third-generation cephalosporins and cotrimoxazole. It belonged to the AFR13 sublineage and clustered with *V. cholerae* isolates collected in Yemen from 2016 to 2019. This second Lebanese strain also harboured the same multidrug-resistance (MDR) IncC-type plasmid found in Yemeni isolates from 2018.

**Interpretation:** The 2022-2023 Lebanese cholera outbreak was caused by the simultaneous introduction of two different 7PET strains. The MDR strain was geographically limited, but the spread of this clone or the horizontal transfer of the MDR plasmid to more susceptible clones could affect epidemic cholera case management. Genomic surveillance is crucial to prevent further spread, and to ensure a prompt and effective response to outbreaks.

**Funding:** The study was funded by the Centers for Disease Control (CDC) award number BAA 75D301-21-C-12132, a grant awarded to the American University of Beirut, WHO country office Lebanon, the Lebanese University, and Institut Pasteur.

**RESEARCH IN CONTEXT PANEL:** *Evidence before this study:* Whole-genome sequencing (WGS) has greatly advanced our understanding and the characterisation of *Vibrio cholerae* outbreaks. However, few studies in the Middle East and North Africa (MENA) region have used this powerful technology. We searched PubMed for studies investigating the molecular epidemiology of *V. cholerae* by WGS in the MENA region, including Lebanon, with the terms “cholera*” AND “a country name of MENA countries” with no restrictions on language or date. The very small number of studies identified concerned Yemen and Algeria. All the outbreaks in the MENA region investigated to date and many others worldwide were caused by a single strain introduced once, contrasting with the endemic setting (the Bay of Bengal) in which several lineages circulate together. One manuscript addressing the history of cholera in Africa from a genomic perspective assigned three Lebanese strains from past outbreaks in 1970 and 1993 as O1 Ogawa isolates from waves 1 and 2 of the seventh pandemic lineage (7PET).

*Added value of the study:* We provide the first comprehensive overview of the molecular epidemiology of the *V. cholerae* strains responsible for the 2022-2023 Lebanese cholera outbreak. The use of WGS made it possible to distinguish clearly between two phylogenetically distant strains from genomic wave 3 of the 7PET lineage responsible for the Lebanese outbreak and to assign their putative origins to South Asia and Yemen. Based on their different susceptibility patterns (a predominant strain with a narrow resistance profile and a minor strain with an extended resistance profile), WGS excluded the hypothesis of the multidrug-resistant (MDR) minor strain emerging from the susceptible dominant strain through the acquisition of the MDR plasmid, instead clearly demonstrating the seeding of the outbreak by two different introductions.

*Implications of all available evidence:* This study demonstrates the importance of WGS associated with national surveillance for obtaining new insights and perspectives, modifying our perception of *V. cholerae* outbreak. This unexpected occurrence of a two-strain outbreak in a setting considered non-endemic for *V. cholerae* requires tight control by the local health authorities to prevent the sporadic introduction and spread of additional strains. Our findings raise the question of the extent to which the strains identified, particularly those from South Asia, spread in Iraq and Syria, neighbouring countries that declared cholera outbreaks before Lebanon. It is difficult to answer this question due to the lack of strains collected from these countries. Regional surveillance of the causal agent of cholera is therefore essential, to unravel transmission events and monitor the emergence of antimicrobial drug-resistant strains observed in many countries around the world.

## INTRODUCTION

Cholera is an acute life-threatening diarrhoeal disease caused by two cholera toxin-producing serogroups — O1 and, less frequently, O139 — of a Gram-negative bacterium, *Vibrio cholerae;* it occurs following the ingestion of contaminated water and food in endemic and epidemic settings.^1^ Cholera continues to be a major public health problem, with 1.3 to 4.0 million cases and 21,000 to 143,000 deaths annually according to the World Health Organisation (WHO).^2^ Cholera outbreaks are still raging globally, particularly in countries already bearing the brunt of natural disasters, human turmoil, and weak economic systems. Indeed, the two countries hardest hit by outbreaks in modern history are Yemen (2016-present) and Haiti (2010-2019; September 2022-present)) with record numbers of cases, at 2.5 million (as recorded in April 2021) and 820,000, respectively.^3,4^

Lebanon has witnessed several cholera outbreaks throughout its history, the most recent of these past outbreaks occurring between July and December 1993, with a total of 344 cases and 29 deaths.^5^ On October 6^th^, 2022, the Lebanese Ministry of Public Health (MoPH) notified the WHO of two laboratory-confirmed cases of cholera meeting all diagnostic criteria in the absence of an epidemiological link with a confirmed cholera outbreak (i.e. culture, including seroagglutination with *V. cholerae* O1-specific antisera, and confirmation of the presence of the cholera toxin genes by PCR) reported by the North and Akkar governorates in northern Lebanon. The index case, a 51-year-old Syrian man living in an informal settlement in Minieh-Donniyeh district (North governorate), was reported on October 5^th^, 2022. This patient was admitted to hospital on the October 1^st^, with rice-water diarrhoea and severe dehydration. The second case occurred in a 47-year-old health worker, possibly through healthcare-associated transmission and corresponding to the first nosocomial infection of this outbreak. Shortly after these two cases were identified, the epidemiological surveillance unit began detecting active cases in the informal settlement inhabited by the index case. In total, about ten additional cases were diagnosed by laboratory testing. *V. cholerae* was also found in sources of drinking water, irrigation, and sewage (October 9, 2022). In parallel, two culture-confirmed cases were identified in Halba (the capital of Akkar Governorate). On the October 10^th^, an additional four cases were confirmed by culture in Syrian nationals living in an informal settlement in Aarsal, a town in the Baalbek district.

Within three months, the cholera outbreak spread across all eight governorates and 20 of the 25 districts in Lebanon (Figure 1). The last positive case was recorded on January 5^th^, 2023. As of June 2023, the cumulative number of reported cholera cases was 8,007, with 671 cumulative culture-confirmed cases and 23 deaths. The outbreak was concentrated in the northern governorates of Akkar and North Lebanon, and in the Beqaa Valley. About 29% of the cases concerned children aged 0-4 years, and 16% of all cases have required hospitalisation. Daily hospitalisation rates peaked one week into the outbreak, with more than 220 patients hospitalised per day. During this period, the case fatality rate (CFR) for cholera reached 11% and the attack rate was highest in the northern districts of Akkar and Minieh-Donniyeh. The intensity of the outbreak necessitated the activation of a multisectoral response to increase cholera preparedness, including greater laboratory capacity for cases of acute watery diarrhoea (AWD) (testing of stools and water) and the training of surveillance and rapid response teams in the early identification of AWD cases. The response to cholera was also strengthened by introducing the oral cholera vaccine (OCV) and ensuring the maintenance of adequate support for intensive care units to prepare for emergencies and the provision of training in the management of cholera and other forms of AWD. The massive cholera vaccination campaign (more than 1 million people) conducted up to January 2023 helped to contain the disease but did not entirely eliminate the risk, particularly as only a single dose of vaccine was administered, rather than the recommended two doses, thus providing protection against cholera for about six months, rather than two years. There is therefore a risk of a resurgence of the disease.

**Figure 1.**
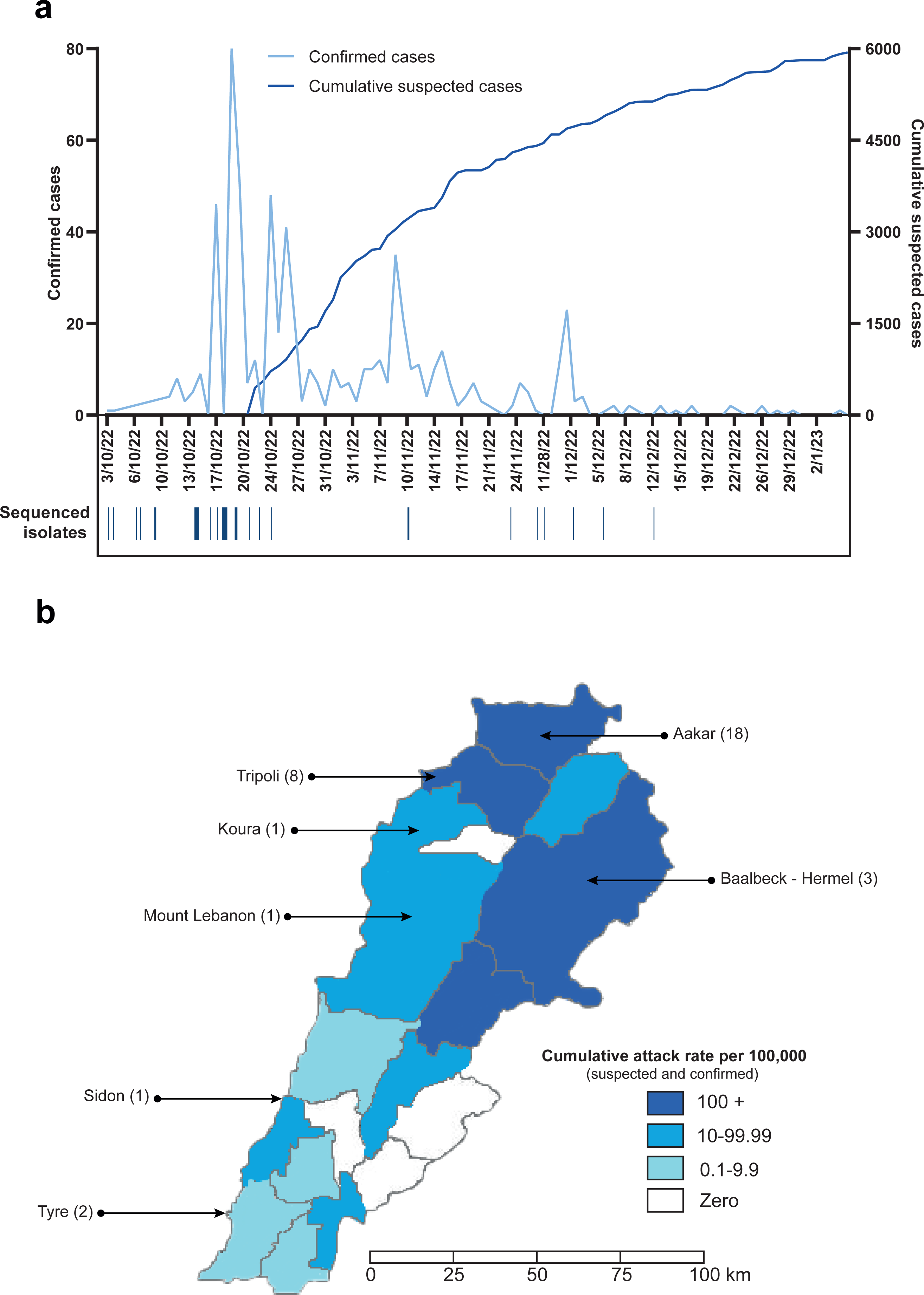
Geographic location at which the *V. cholerae* O1 El Tor isolates sequenced were obtained and number of reported cholera cases. a, Cumulative number of suspected cholera cases per day vs. the confirmed number of cases per day in Lebanon until January 3, 2023. The dates on which the isolates sequenced in this study were obtained are shown under the epidemic curve. b, Geographic locations at which the 34 sequenced *V. cholerae* O1 El Tor isolates were obtained in Lebanon.

Two different antimicrobial drug resistance (AMR) profiles differing in prevalence were observed in the *V. cholerae* O1 isolates obtained during the 2022-2023 Lebanese outbreak: a major profile with a narrower resistance spectrum and a minor profile with an extended multidrug-resistant (MDR) profile. Two scenarios could potentially account for this distribution: the first, considered the most likely *a priori*, involves the acquisition of a mobile genetic element, such as an MDR plasmid, by the strain with a narrower resistance profile initially responsible for the outbreak, following some kind of selection pressure and thereby resulting in an MDR profile (as in Yemen). ^6^ The second scenario, which is uncommon outside the Bay of Bengal, involves the circulation of two different strains introduced separately into Lebanon. We used whole-genome sequencing (WGS) and genomic epidemiology approaches to determine which of these two scenarios had occurred in Lebanon. We also delved into the genetic basis of antibiotic resistance and virulence in the circulating isolates.

## MATERIALS AND METHODS

### Ethics statement

This research study was based exclusively on bacterial isolates and the corresponding metadata collected for nationwide surveillance of the cholera outbreak by the Lebanese Ministry of Public Health (MoPH) in collaboration with the American University of Beirut (AUB), Rafic Hariri University Hospital, and the “Laboratoire Microbiologie Santé et Environnement” (LMSE). Hence, neither informed consent nor institutional review board (IRB) approval was required.

### Vibrio cholerae isolates

Stools, sewage, water and plant samples were collected by the MoPH and delivered to the bacteriology and molecular microbiology research laboratory at AUB, a WHO collaborating centre for reference and research on bacterial pathogens. In total, 671 clinical isolates of *V. cholerae* were identified, with 144 isolates from North Lebanon collected and stored in the “la Collection Microbiologique de l’Université Libanaise (CMUL)” at LMSE at the Lebanese University. We included 18 isolates from AUB and 16 from the LMSE in this study (Appendix 1). These isolates were recovered between October and December 2022, for continuous surveillance and prevention, the last positive case being reported on January 5^th^, 2023.

### Bacterial culture and identification

The approach to bacterial culture and identification differed between the LMSE and AUB laboratories. At the LMSE, part of each stool sample was plated directly on two different media, a non-selective nutrient-rich agar medium (pH=8.5, 10 g/l NaCl), and a selective agar medium, thiosulphate-citrate-bile salts-sucrose (TCBS) agar (BioMérieux, Marcy-l’Etoile, France). Another portion of each sample was incubated in alkaline peptone water (Bio-Rad, Marnes-la-Coquette, France; 10 g/l NaCl) for 6-8 hours at 35-37°C and was then plated on the same solid media. By contrast, at AUB, the stool sample was incubated in alkaline peptone water for 6 hours at 35°C and was then plated on the surface of TCBS agar, MacConkey agar (Bio-Rad), and *Vibrio* Chromagar (CHROMagar, Paris, France). After standard microbiological identification by microscopy (a comma-shaped Gram-negative bacterium) and oxidase tests, *V. cholerae* isolates were identified with API 20E test strips (BioMérieux) and by matrix-assisted laser desorption ionisation–time of flight mass spectrometry (MALDI-TOF), with Vitek MS (BioMérieux) at LMSE or MALDI Biotyper (Bruker Daltonics, Germany) at AUB. Agglutination was performed with specific antisera (Artron Laboratories Inc., British Columbia, Canada for O1 and O139 antisera) and the presence of cholera toxin genes was confirmed with a multiplex PCR method developed by Hoshino and colleagues,^7^ for the first five cases.

### Antimicrobial drug susceptibility testing

Antimicrobial drug susceptibility testing was performed by the disk diffusion method in accordance with Clinical and Laboratory Standards Institute (CLSI) guidelines. Tests were performed for 14 antimicrobial agents: piperacillin/tazobactam, cefotaxime, ceftazidime, meropenem, nalidixic acid, ciprofloxacin, levofloxacin, erythromycin, azithromycin, trimethoprim-sulfamethoxazole (cotrimoxazole), tetracycline, doxycycline, nitrofurantoin, and vibriostatic agent O/129. CLSI interpretative criteria for the antibiotic susceptibility testing of *Vibrio spp*. (M45 document) were used when available.^8^ For vibriostatic agent O/129 (equivalent to trimethoprim), nitrofurantoin, nalidixic acid, and ciprofloxacin, the interpretative criteria for *Enterobacteriaceae*/*Salmonella* spp. (M100-S30 document) were used.^9^ The minimum inhibitory concentrations (MICs) of colistin for 19 isolates — 16 from the LMSE and 3 from the AUB — were determined at Institut Pasteur with the SensititreTM system (Thermo Fisher Scientific).

### Whole-genome sequencing

We studied 34 *V. cholerae* isolates, 16 originating from the LMSE, which were sequenced at the Institut Pasteur in Paris, and 18 isolates originating from AUB, which were sequenced in-house. At Institut Pasteur, genomic DNA was extracted with the Maxwell 16-cell purification kit (Promega, https://www.promega.com). The DNA libraries were then prepared at the Institut Pasteur Mutualized Platform for Microbiology (P2M) with the Nextera XT kit (Illumina, San Diego, CA, USA) and sequencing was performed with the NextSeq 500 system (Illumina), generating 150 bp paired-end reads. At AUB, genomic DNA was extracted with the Quick-DNA™ Fungal/Bacterial Miniprep kit (Zymo Research, Irvine, CA) and purified with the Genomic DNA Clean & Concentrator™ kit (Zymo Research), according to the manufacturer’s protocols. DNA libraries were prepared with the Illumina DNA prep kit (Illumina GmbH, Munich, Germany) and subjected to Illumina MiSeq 2 × 150 bp paired-end sequencing by Illumina MiSeq.

The resulting reads were filtered with FqCleanER version 21.10 (https://gitlab.pasteur.fr/GIPhy/fqCleanER) with options -q 28 -l 70 to discard low-quality reads with phred scores <28 and length <70 bp and to remove adaptor sequences.^10^ Two isolates were selected for long-read sequencing, based on geographic origin and representativity of the two circulating strains. Long-read sequencing was performed on isolate CNRVC220127 (alternative name CMUL 009) at Institut Pasteur, with a MinION nanopore sequencer (Oxford Nanopore Technologies), as previously described.^6^ The second isolate sequenced (VIC_202210_72, alternative name VIC11A) was cultured on MacConkey agar. DNA was extracted with the Quick DNA Fungal/Bacterial Miniprep Kit (by ZYMO Research) and cleaned with DNA Clean & Concentrator -5 (by ZYMO Research). The clean DNA was then sequenced with an Oxford Nanopore MinION at AUB. The DNA library was prepared with Rapid Barcoding Kit 96 (SQK-RBK110.96) and sequenced on R9.4.1 flow cells (FLO-MIN106).

The sequences of the two isolates were assembled from both long and short reads, by two different methods. For CNRVC220127, a hybrid approach was used in UniCycler v.0.4.8.^10^ For VIC_202210_72, a combination of Raven^11^ v.1.6.0 (https://github.com/lbcb-sci/raven), Medaka v.1.4.4 (https://github.com/nanoporetech/medaka) and Polypolish^12^ v.0.5.0 (https://github.com/rrwick/Polypolish/) was used. The large plasmid of VIC_202210_72 was then annotated with Bakta^13^ v.1.5.0, corrected manually and visualised with BRIG v.0.95 (http://sourceforge.net/projects/brig).^10^

### Genomic analysis

For construction of a globally representative set of isolates, we downloaded and included in this study sequences available either in raw-read format or as assembled genomes in the European Nucleotide Archive (ENA) (https://www.ebi.ac.uk/ena) and GenBank (https://www.ncbi.nlm.nih.gov/genbank/) databases (Appendix 2).

The phylogenomic analysis was performed as previously described.^14^ Briefly, the paired-end reads were mapped onto the reference genome of *V. cholerae* O1 El Tor strain N16961, also known as A19 (GenBank accession numbers LT907989 and LT907990) with Snippy. The single-nucleotide variants (SNVs) were then called with Snippy v.4.6.0/Freebayes v.1.3.2 (https://github.com/tseemann/snippy), using the following parameters: a minimum read coverage of 4, a minimum base quality of 13, a mapping quality of 60, and a 75% read concordance at a locus for a variant to be reported. Finally, core-genome SNVs were then aligned in Snippy for phylogenetic inference.

Repetitive sequences (insertion sequences and the TLC-RS1-CTX region) and recombinogenic regions (VSP-II) were masked.^15^ Putative recombinogenic regions were identified and masked with Gubbins v.3.2.0.^10^ A maximum likelihood (ML) phylogenetic tree was constructed from an alignment of 10,632 chromosomal SNVs, with RAxML v. 8.2.12, under the GTR model, with 200 repetitions for bootstrapping.^16^ This global tree was rooted on the A6 genome and visualised with iTOL v.6 (https://itol.embl.de/).^17^

Short reads from Illumina were assembled *de novo* with SPAdes v.3.15.2.^10^ The presence of various genetic markers (O1 *rfb* gene, whole locus of VSP-II, *ctxB*, and *wbeT*) was investigated with BLAST v.2.2.26 against reference sequences, as previously described.^14,15^ The presence and type of acquired antibiotic resistance genes (ARGs) or ARG genetic structures were investigated with ResFinder v.4.0.1 (https://cge.cbs.dtu.dk/services/ResFinder/), PlasmidFinder v.2.1.1 (https://cge.cbs.dtu.dk/services/PlasmidFinder/), and BLAST analysis against GI-15, Tn*7*, and SXT/R391 integrative and conjugative elements (ICE).^15^ The sequences assembled *de novo* were examined with BLAST to look for mutations of genes encoding resistance to nitrofurans (*VC_0715* and *VC_A0637*), resistance to quinolones (*gyrA* and *parC*) or restoring susceptibility to polymyxin B (*vprA*), as previously described.^15,18^

### Data availability

Short reads were submitted to the ENA under study project PRJEB65303 (Appendix 2). Assemblies resulting from long-read sequencing were submitted to GenBank under project PRJNA1013428 (CP134060-CP134061 for CNRVC220127/CMUL009 and CP134057-CP134059 for VIC11-A).

## RESULTS

### Antimicrobial drug susceptibility testing results

Antimicrobial drug susceptibility testing in the Lebanese laboratories (LMSE and AUB) identified two different AMR profiles (Table 1 and Appendix 1) in the 671 *Vibrio cholerae* O1 isolates recovered during the outbreak. One of these profiles predominated, accounting for 94.7% (636/671) of the isolates collected across Lebanon, including the 19 isolates tested at Institut Pasteur. It displayed resistance to nitrofurantoin, the vibriostatic agent O/129, and nalidixic acid only, and decreased susceptibility to ciprofloxacin. The second profile was a minor profile accounting for only 5.6% (35/671) of the isolates. It was characterised by resistance to third-generation cephalosporins (cefotaxime and ceftazidime), macrolides (erythromycin and azithromycin), sulphonamides, the vibriostatic agent O/129, cotrimoxazole, nitrofurantoin and nalidixic acid, and decreased susceptibility to ciprofloxacin. This second AMR profile was found exclusively in isolates originating from the South of Lebanon (Tyr) and Beqaa.

**Table 1.** Characteristics of the two epidemic strains of *Vibrio cholerae* O1 involved in the cholera outbreak in Lebanon in 2022-2023*.

| Category | Predominant strain (*n* = 31) | Minor strain (*n* = 3) |
| --- | --- | --- |
| Serogroup and serotype, | O1, Ogawa | O1, Ogawa |
| Sequence type | ST69 | ST69 |
| Lineage | 7PET | 7PET |
| Genomic wave | 3 | 3 |
| Genetic markers | *ctxB7*, *tcpA*<sup>CIRS101</sup>, VSP-IIΔ | *ctxB7*, *tcpA*<sup>CIRS101</sup>, VSP-IIΔ |
| AMR profile, selected antimicrobial drugs |  |  |
| Cefotaxime | Susceptible | Resistant |
| Meropenem | Susceptible | Susceptible |
| Erythromycin | Susceptible | Resistant |
| Azithromycin | Susceptible | Resistant |
| Nalidixic acid | Resistant | Resistant |
| Ciprofloxacin | Intermediate | Intermediate |
| Tetracycline | Susceptible | Susceptible |
| O/129 | Resistant | Resistant |
| Trimethoprim/<br>sulfamethoxazole | Susceptible | Resistant |
| Nitrofurantoin | Resistant | Resistant |
| Colistin | Susceptible | Susceptible |
| Horizontally acquired AMR elements | ICE*Vch*Ind5Δ<sup>†</sup> | ICE*Vch*Ind5Δ, pCNRVC190243<sup>‡</sup> |
| Horizontally acquired AMR genes | *dfrA1*<sup>‡</sup> | *dfrA1*, *aadA2*, *sul1*, *bla*<sub>PER-7</sub>, *mph(A)*, *msr(E)*, *mph(E)* |

| Chromosomal gene mutations | AMR phenotype | AMR phenotype |
| --- | --- | --- |
| *gyrA*_S83I and *parC*_S85L | Resistance to nalidixic acid; decreased susceptibility to ciprofloxacin | Resistance to nalidixic acid; decreased susceptibility to ciprofloxacin |
| *VC0715*_R169C and *VCA0637*_Q5Stop | Resistance to nitrofurantoin | Resistance to nitrofurantoin |
| *vprA*_D89N | Susceptibility to colistin | Susceptibility to colistin |

AMR, antimicrobial resistance; 7PET, seventh pandemic *V. cholerae* biotype El Tor lineage; VSP-IIΔ, deletion encompassing VC0495–VC0512 (according to GenBank accession no. AE003852) in *Vibrio* seventh pandemic island II (VSP-II); ICE*Vch*Ind5Δ, deletion encompassing ICE*Vch*Ind50011–ICE*Vch*Ind50019 (according to GenBank accession no. GQ463142) in ICE*Vch*Ind5, an integrative conjugative element (ICE) of the SXT/R391 family; †one isolate (VIC\_202211\_60) also had a Col3M plasmid carrying the *qnrD1* gene,

### Phylogenetic and genomic features of *V. cholerae* O1 isolates

We selected 34 *V. cholerae* O1 isolates for further analysis on the basis of their AMR profiles (Figure 1a); 31 presented the predominant AMR profile (limited resistance) and three presented the minor AMR profile (extended resistance) (Table 1). Genome sequencing confirmed that all 34 isolates belonged to serogroup O1, serotype Ogawa and biotype El Tor (sequence type ST69). All the *V. cholerae* O1 isolates (Table 1) displayed the following genomic features: (i) the *ctxB7* variant of the cholera toxin subunit B gene, (ii) the toxin-coregulated pilus gene subunit A gene variant *tcpA*^CIRS101^, and (iii) a deletion (ΔVC0495–0512) in the *Vibrio* seventh pandemic island II (VSP-II) (Table 1).

All 34 *V. cholerae* O1 isolates had (i) a deletion of about 10 kb in the chromosomal ICE*Vch*Ind5 integrative and conjugative element, resulting in the loss of four genes encoding resistance to streptomycin (*strA* and *strB*), sulphonamides (*sul2*), and chloramphenicol (*floR*), but not the fifth gene encoding resistance to the vibriostatic agent O/129 (*dfrA1*), (ii) mutations of the chromosomal *VC0715* (resulting in the R169C substitution) and *VCA0637* (resulting in a premature codon stop at Q5) nitroreductase genes, leading to nitrofurantoin resistance, (ii) a mutation of the chromosomal *VC1320* (*vprA*) (D89N) gene re-establishing susceptibility to polymyxin B, and (iv) mutations of the chromosomal DNA gyrase *gyrA* (S83I) and topoisomerase IV *parC* (S85L) genes, leading to nalidixic acid resistance and decreased susceptibility to ciprofloxacin. The three isolates with the second AMR profile also carried an IncC of about 139 kb (formerly IncA/C_2_). This plasmid displayed 100% nucleotide sequence identity to the MDR plasmid, pCNRVC190243 (GenBank accession number OW443149.1),^6^ found in *Vibrio cholerae* O1 isolates from Yemen in 2018-2019 (Figure 2).^6^ The backbone of pCNRVC190243 harboured a 20 kb pseudo-compound transposon, Yem*Vch*MDRI, flanked by IS*26* insertion sequences and encompassing genes encoding aminoglycoside resistance (*aadA2*), a quaternary ammonium compound efflux pump (*qac*), sulphonamide resistance (two copies of the *sul1* gene), an extended-spectrum beta-lactamase (ESBL; *bla*_PER-7_), and macrolide resistance (*mph(A)*, *mph(E)*, and *msr(E)*). The resistance to cotrimoxazole of isolates with the second AMR profile probably resulted from the simultaneous presence of the plasmid-borne *sul1* gene and the chromosomal *dfrA1* gene. The pCNRVC190243 plasmid had a nucleotide sequence 99.98% identical to that of pYA00120881 (GenBank accession number MT151380) identified in Zimbabwean *Vibrio cholerae* O1 isolates collected in 2015 and 2018, but it carried a different multidrug-resistance region, containing, in particular, the ESBL gene *bla*_CTX-M-15_.^19^

**Figure 2.**
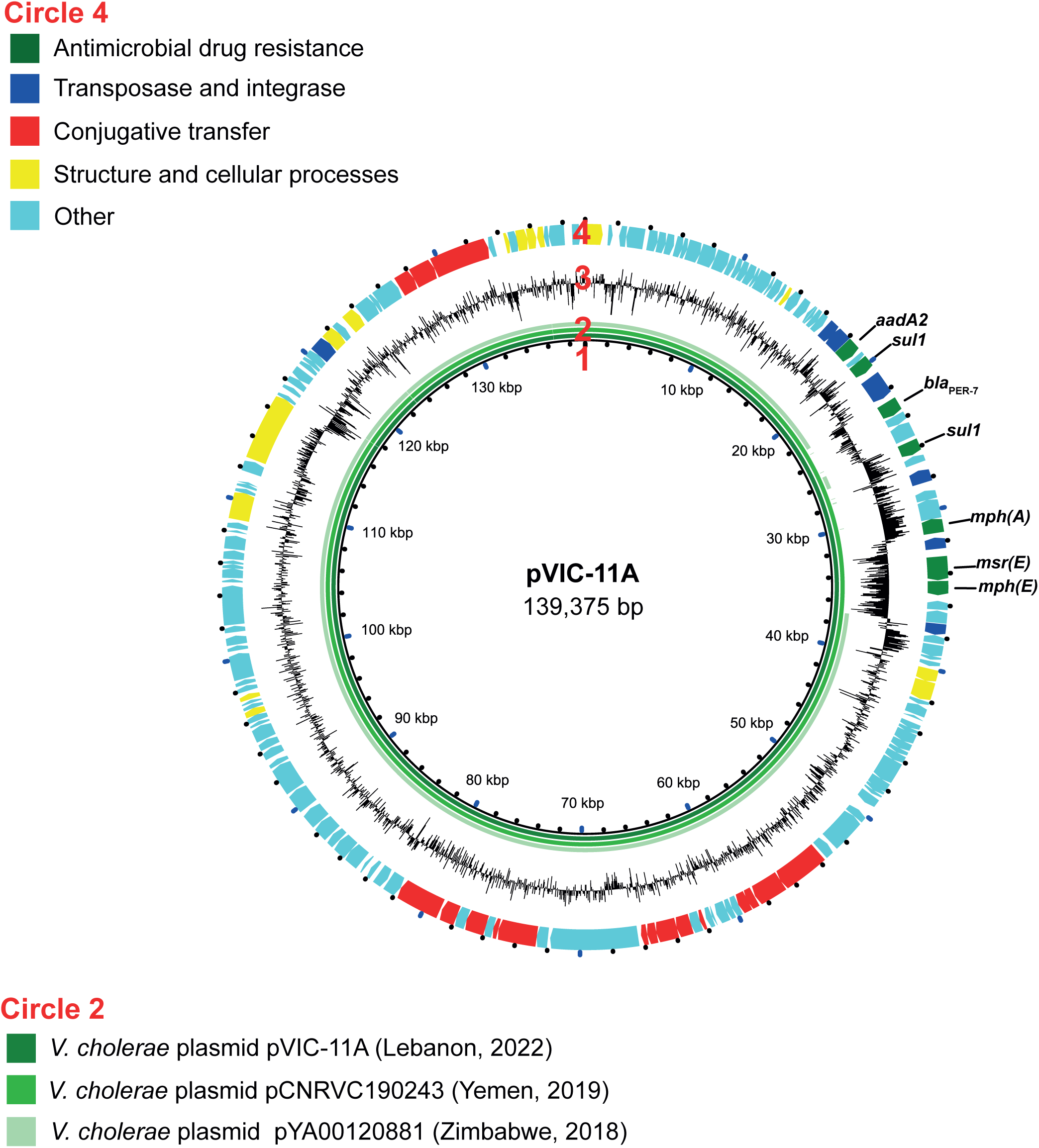
Circular map and comparative analysis of the IncC2 plasmid found in some *V. cholerae* O1 isolates from Lebanon in 2022. Circles from innermost to outermost indicate (1) the nucleotide position of the plasmid of the VIC_202210_72 isolate (alternative name pVIC-11A), (2) the alignment throughout the plasmids between pVIC-11A (Lebanon, 2022) in dark green, pCNRVC190243 (Yemen, 2019) (GenBank accession number OW443149.1) in medium green, and pYA00120881 (Zimbabwe, 2018) (GenBank accession number MT151380) in light green, (3) the G+C content map of pVIC-11A, and (4) the coding sequences (CDS) map of pVIC-11A, in which green arrows indicate antimicrobial drug resistance CDS, dark blue arrows transposase and integrase CDS, red arrows the CDS involved in conjugative transfer, yellow arrows those involved in the structure and cellular processes, and light blue CDS with other functions. The names of resistance genes within the YemVchMDRI are indicated above the corresponding CDS.

One isolate (VIC_202211_60) also harboured a putative small Col3M colicin plasmid encoding a plasmid-mediated quinolone resistance protein, QnrD1, a member of the Qnr family protecting DNA-gyrase and topoisomerase IV against quinolones. However, this had no effect on the quinolone resistance profile of this isolate.

We then placed these 34 *V. cholerae* O1 isolates from the Lebanon 2022-2023 outbreak in a global context by constructing a maximum-likelihood phylogeny of 1,465 7PET genomic sequences using 10,632 SNVs evenly distributed over the non-repetitive, non-recombinant core genome. All 34 isolates clustered in the genomic wave 3 clade of the 7PET lineage, and more particularly in the subclade containing isolates with the *ctxB7* allele (Figure 3). These 34 isolates also differed from other isolates previously recovered in Lebanon, including those isolated in 1970 and 1993, which clustered within genomic waves 1 and 2 of the 7PET lineage, respectively (Figure 3).

**Figure 3.**
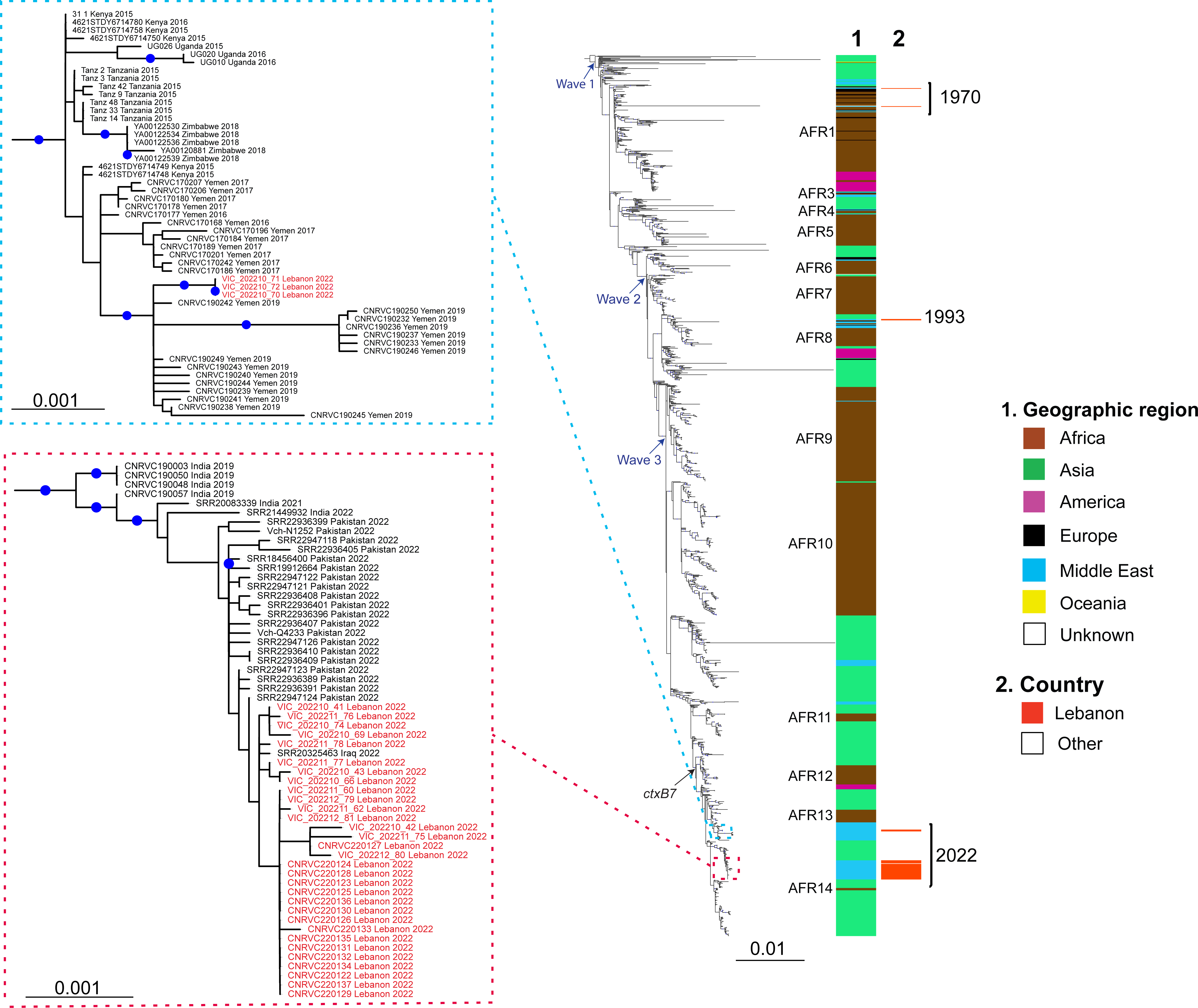
Maximum-likelihood phylogeny of *Vibrio cholerae* O1 El Tor isolates collected in Lebanon in 2022, compared with 1,465 reference seventh pandemic *V. cholerae* El Tor genomic sequences. A6 was used as the outgroup. Blue arrows represent the three genomic waves and the black arrow indicates the acquisition of the *ctxB7* allele. The colour coding in the first column shows the geographic origins of the isolates, and African sublineages (AFR1, AFR3–AFR14) are shown on the left. The red colour in the second column indicates the Lebanese origin of the isolates. A magnification of the clades containing the two strains from Lebanon (red square corresponding to the predominant strain and blue square corresponding to the multidrug-resistant minor strain) is shown on the left with red text indicating the Lebanese isolates. For each genome, its name (or accession number), the country in which contamination occurred and the year of sample collection are indicated at the tip of the branch. Scale bars indicate the number of nucleotide substitutions per variable site. Blue dots correspond to bootstrap values ≥90%.

Our phylogenomic analysis revealed that the 34 *V. cholerae* O1 isolates were distributed between two different clusters according to their AMR profiles (Figure 3). Indeed, the 31 isolates with the predominant profile clustered together (median pairwise distance of 1.5 [range 0–9] core-genome SNVs) and with many other isolates originating from the Pakistan 2022 outbreak, from India (2019-2022), and one isolate from the Iraq 2022 outbreak. The three remaining isolates with the extended AMR profile clustered together (no SNV between them) and with Yemeni isolates recovered between 2016 and 2019.

## DISCUSSION

After approximately three decades without cholera, Lebanon recently suffered an outbreak extending from October 2022 to January 2023 (Figure 1).^20^ The two different AMR profiles observed in the Lebanese isolates initially suggested the possibility of a mother strain relatively susceptible to antibiotics acquiring an MDR plasmid early in the outbreak, or of two different strains circulating simultaneously, this second possibility being considered less likely. However, the high discrimination power of WGS made it possible to distinguish two different strains of *V. cholerae* O1 serotype Ogawa harbouring the *ctxB7* allele from genomic wave 3 of the 7PET lineage, and, thus, to conclude that the 2022-2023 cholera outbreak in Lebanon was caused by two phylogenetically distant strains rather than a single strain that subsequently acquired an MDR plasmid. The two-strain outbreak scenario was initially considered unlikely because outbreaks in countries non-endemic for cholera generally occur following a single introduction of a single strain, contrasting with the situation in the Bay of Bengal, where many lineages circulate simultaneously.^21,22^ This two-strain outbreak in Lebanon is thus unusual, as it stems from two different introductions outside the endemic setting. The two strains concerned had different AMR profiles and different patterns of circulation in Lebanon. The strain with the narrower AMR profile predominated in all affected regions of Lebanon, including North Lebanon in particular, whereas the strain with the broader AMR profile was found only in South Lebanon and Beqaa.

Globally, the 7PET lineage has been responsible for the repeated spread of the seventh pandemic from the Bay of Bengal in South Asia to the rest of the world through three epidemic waves.^15,23^ The wave 3 clade carrying the *ctxB7* allele first emerged in Kolkata (India) in 2006,^24^ subsequently spreading to other parts of the world, including Haiti and Yemen, and across Africa.^6,19,25,26^

In the global phylogenetic tree, the predominant Lebanese strain clustered with isolates from South Asia, including isolates from the 2022 Pakistani outbreak collected locally (GenBank bioproject PRJNA916827, https://www.ncbi.nlm.nih.gov/bioproject/?term=PRJNA916827) or in travellers from the US and Australia with a history of travel to Pakistan.^14^ These isolates were considered the direct ancestors of the Lebanese strain (Figure 3). Two very recently published studies revealed that the strain circulating in the 2022 Pakistani outbreak was also the ancestor of a strain circulating in South Africa and Malawi considered to belong to the AFR15 sublineage.^14,27^ Indeed, although the Lebanese strains were not identified and analysed in the studies performed in Malawi, their phylogenetic tree incorporated the same genomes from Pakistan and showed the grouping of these genomes with the AFR15 sublineage, thereby revealing similarity between the predominant Lebanese strain and the AFR15 sublineage. A sublineage closely related to AFR15 may therefore have been imported into the Middle East region directly from South Asia or indirectly via Africa. Interestingly, one isolate (PNUSAV00294, SRR20325463)^14^ collected in the US in June 2022 from a traveller with a history of recent travel to Iraq clustered with the Lebanese isolates, suggesting that the predominant strain in this Lebanese outbreak was the same strain that swept the region (Iraq and Syria) shortly before the Lebanese outbreak, causing outbreaks beginning on June 20^th^, 2022 in Iraq, ^28^ and September 10^th^, 2022 in Syria.^29^ One month after the declaration of the Syrian outbreak, Lebanon declared its first index case in a Syrian refugee residing in North Lebanon, providing additional support for the theory that the predominant strain in Lebanon was imported from Syria. According to the United Nations High Commission for Refugees, Lebanon has the largest refugee population per capita in the world, with an estimated 1.5 million Syrian refugees living on its soil. Nevertheless, our ability to infer precise transmission routes through genomic analysis is hampered by the lack of availability of isolates from the various affected countries in the region, including Syria and Iraq.

The minor strain displaying MDR grouped with isolates from the 2019 outbreak in Yemen, suggesting the probable direct introduction of this strain from Yemen into South Lebanon and Beqaa, the areas from which it was exclusively isolated. Like the Yemeni isolates, this strain carried an IncC plasmid (pCNRVC190243) bearing determinants of resistance to cotrimoxazole, macrolides, third-generation cephalosporins and aminoglycosides. The Yemeni epidemic comprised several waves but was seeded by a single introduction linked to the 7PET sublineage AFR13, which was recently transmitted from South Asia into East Africa and from there to Yemen.^6,18^ Before 2018-2019, AFR13 isolates had features resembling those of the predominant Lebanese strain, including a narrow resistance profile, due partly to an ICE*Vch*Ind5 deletion and the subsequent loss of four of five AMR genes. However, a plasmid-carrying AFR13 clone, the pCNRVC190243-carrying AFR13 clone, began to emerge in late 2018 and displayed resistance to many therapeutically relevant drugs. The spread of this MDR clone was driven by the therapeutic overuse of macrolides.^6^ Our multidrug-resistant minor strain is, therefore, a Yemeni AFR13 clone carrying a self-transmissible MDR plasmid. The spread of this strain would greatly decrease treatment options and jeopardise cholera case management. This bleak scenario might occur if the multidrug-resistant Yemeni clone manages to expand beyond its current geographic location and spread throughout Lebanon or if it passes its plasmid to the predominant strain (sublineage closely related to AFR15).

Similar IncC plasmids were previously observed in other *V. cholerae* strains, including the pYAM00120881 plasmid identified in Zimbabwe in 2018, which has a backbone almost identical to that of pCNRVC190243.^19^ Intriguingly, the pYAM00120881-carrying AFR13 clone caused a six-month-long outbreak in Zimbabwe, with over 10,000 suspected cases,^19^ demonstrating a certain degree of plasmid stability in a 7PET *V. cholerae* stain sustaining an outbreak and even providing evidence against the presumed plasmid instability in this species.^6^ It has been suggested that the 10 kb deletion in SXT/R391 ICE (IC*EVch*Ind5) seen in our isolates might render these isolates fit to host MDR IncC plasmids stably, with deleterious consequences for antibiotic susceptibility in the future.^6^

Our genomic analysis revealed general consistency with the phenotypic AMR profile. All the isolates also harboured the *catB9* gene, but this gene does not confer chloramphenicol resistance,^30^ consistent with the chloramphenicol susceptibility of the isolates (data not shown). Resistance to polymyxin B has been used as a marker of *V. cholerae* O1 biotype El Tor since the start of the seventh cholera pandemic, as a means of differentiating this biotype from the classical biotype, which was susceptible to polymyxin B. The susceptibility to polymyxins of our isolates may be due to the VprA (VC1320) D89N substitution, as previously reported in the AFR13 and AFR14 sublineages.^18,26^

Lebanon has been struggling with an unprecedented, multifaceted crisis since 2019, including severe economic collapse, the COVID-19 pandemic, the explosion in the Port of Beirut in August 2020 and a high burden of refugees. This major crisis and the fragile infrastructure of Lebanon favoured this outbreak. The economic crisis, with all its implications, affected all aspects of the outbreak. Access to safe water was hampered by the lack of electricity and the inadequacy of the sewage system. The country was already suffering from a shortage of medical and diagnostic supplies in addition to the global shortage of oral cholera vaccines, and laboratory supplies for cholera diagnosis. Nevertheless, tremendous collaborative efforts were initiated, under the auspices of the Ministry of Public Health in Lebanon and in collaboration with several national and international organisations including, but not limited to the WHO, UNICEF, UNHCR, and ICRC, making it possible to halt the spread of the disease within three months of the declaration of the index case. However, we are well aware of the possibility of disease resurgence and of another outbreak, particularly as the neighbouring countries have not yet brought their own outbreaks under control.

In conclusion, the outbreak in Lebanon was caused by two different strains: one with a narrower AMR profile related to South Asian isolates and the other with an extended AMR profile similar to the Yemeni AFR13 *V. cholerae* strain. However, as isolates from the neighbouring countries are missing from the phylogenetic analysis, it may be difficult to establish a comprehensive history for this outbreak. Regional surveillance of the causal agent of cholera by microbial genomics methods is, thus, paramount for the reliable inference of transmission routes and for tracking and monitoring the emergence of any AMR, particularly after the worrying switch of several AFR13 strains from a limited to an extended MDR phenotype following the acquisition of IncC-type plasmids. ^6^

## CONFLICTS OF INTEREST

The authors declare that there are no conflicts of interest.

## ACKNOWLEDGMENT

We would like to thank all the staff and personnel of the laboratories participating in this study, including the American University of Beirut and the American University of Beirut Medical Center, the LMSE, Rafik Hariri University Hospital, the Ministry of Public Health Lebanon, The Lebanese Red Cross, Institut Pasteur, the WHO, and the CDC.

## Notes

### Competing Interest Statement

The authors have declared no competing interest.

## REFERENCES

1 Clemens JD, Nair GB, Ahmed T, Qadri F, Holmgren J. Cholera. The Lancet 2017; 390: 1539–49.

2 Ganesan D, Gupta S Sen, Legros D. Cholera surveillance and estimation of burden of cholera. Vaccine 2020; 38: A13–A17.

3 World Health Organization. CHOLERA SITUATION IN YEMEN. WHO-EM/CSR/434/E. 2021. https://applications.emro.who.int/docs/WHOEMCSR434E-eng.pdf?ua=1

4 World Health Organization. Disease Outbreak News; Cholera – Haiti. 2022. https://www.who.int/emergencies/disease-outbreak-news/item/2022-DON427

5 Harb H. Compiled Literature Report on Selected Health Conditions in Lebanon. 2004. https://www.moph.gov.lb/userfiles/files/Statistics/LiteratureReviewreportonselecteddiseasesinLebanon.pdf

6 Lassalle F, Al-Shalali S, Al-Hakimi M, et al. Genomic epidemiology reveals multidrug resistant plasmid spread between *Vibrio cholerae* lineages in Yemen. Nat Microbiol 2023; 8: 1787–98.

7 Hoshino K, Yamasaki S, Mukhopadhyay AK, et al. Development and evaluation of a multiplex PCR assay for rapid detection of toxigenic *Vibrio cholerae* O1 and O139. FEMS Immunol Med Microbiol 1998; 20: 201–7.

8 CLSI M45-ED3. Methods for Antimicrobial Dilution and Disk Susceptibility Testing of Infrequently Isolated or Fastidious Bacteria. 2016.

9 CLSI M100-S30. Performance Standards for Antimicrobial Susceptibility Testing. 2020.

10 Lefèvre S, Njamkepo E, Feldman S, et al. Rapid emergence of extensively drug-resistant Shigella sonnei in France. Nat Commun 2023; 14: 462.

11 Vaser R, Šikić M. Time- and memory-efficient genome assembly with Raven. Nat Comput Sci 2021; 1: 332–6.

12 Wick RR, Holt KE. Polypolish: Short-read polishing of long-read bacterial genome assemblies. PLoS Comput Biol 2022; 18: e1009802.

13 Schwengers O, Jelonek L, Dieckmann MA, Beyvers S, Blom J, Goesmann A. Bakta: rapid and standardized annotation of bacterial genomes via alignment-free sequence identification. Microb Genom 2021; 7. DOI:10.1099/mgen.0.000685.

14 Smith AM, Sekwadi P, Erasmus LK, et al. Imported Cholera Cases, South Africa, 2023. Emerg Infect Dis 2023; 29. DOI:10.3201/eid2908.230750.

15 Weill F-X, Domman D, Njamkepo E, et al. Genomic history of the seventh pandemic of cholera in Africa. Science *(1979)* 2017; 358: 785–9.

16 Stamatakis A. RAxML-VI-HPC: maximum likelihood-based phylogenetic analyses with thousands of taxa and mixed models. Bioinformatics 2006; 22: 2688–90.

17 Letunic I, Bork P. Interactive Tree Of Life (iTOL) v4: recent updates and new developments. Nucleic Acids Res 2019; 47: W256–W259.

18 Weill F-X, Domman D, Njamkepo E, et al. Genomic insights into the 2016–2017 cholera epidemic in Yemen. Nature 2019; 565: 230–3.

19 Mashe T, Domman D, Tarupiwa A, et al. Highly Resistant Cholera Outbreak Strain in Zimbabwe. New England Journal of Medicine 2020; 383: 687–9.

20 Lebanese Ministry of Public Health. Cholera in Lebanon. 2023. http://www.moph.gov.lb (accessed July 23, 2023).

21 Morita D, Morita M, Alam M, et al. Whole-Genome Analysis of Clinical Vibrio cholerae O1 in Kolkata, India, and Dhaka, Bangladesh, Reveals Two Lineages of Circulating Strains, Indicating Variation in Genomic Attributes. mBio 2020; 11. DOI:10.1128/mBio.01227-20.

22 Mutreja A, Dougan G. Molecular epidemiology and intercontinental spread of cholera. Vaccine 2020; 38: A46–A51.

23 Mutreja A, Kim DW, Thomson NR, et al. Evidence for several waves of global transmission in the seventh cholera pandemic. Nature 2011; 477: 462–5.

24 Naha A, Pazhani GP, Ganguly M, et al. Development and Evaluation of a PCR Assay for Tracking the Emergence and Dissemination of Haitian Variant ctxB in Vibrio cholerae O1 Strains Isolated from Kolkata, India. J Clin Microbiol 2012; 50: 1733–6.

25 Domman D, Quilici M-L, Dorman MJ, et al. Integrated view of Vibrio cholerae in the Americas. Science *(1979)* 2017; 358: 789–93.

26 Benamrouche N, Belkader C, Njamkepo E, et al. Outbreak of Imported Seventh Pandemic Vibrio cholerae O1 El Tor, Algeria, 2018. Emerg Infect Dis 2022; 28. DOI:10.3201/eid2806.212451.

27 Chabuka L, Choga WT, Mavian CN, et al. Genomic epidemiology of the cholera outbreak in Malawi 2022-2023. medRxiv 2023; : 2023.08.22.23294324.

28 Qamar K, Malik UU, Yousuf J, et al. Rise of cholera in Iraq: A rising concern. Annals of Medicine & Surgery 2022; 81. DOI:10.1016/j.amsu.2022.104355.

29 Eneh SC, Admad S, Nazir A, et al. Cholera outbreak in Syria amid humanitarian crisis: the epidemic threat, future health implications, and response strategy – a review. Front Public Health 2023; 11. DOI:10.3389/fpubh.2023.1161936.

30 Kumar P Yadav P NAGAKYPK. Re-emergence of chloramphenicol resistance and associated genetic background in Vibrio cholerae O1. FASEB 2017; 31.

